# PACVr: Plastome Assembly Coverage Visualization in R

**DOI:** 10.1101/697821

**Authors:** Michael Gruenstaeudl, Nils Jenke

## Abstract

**Background:** The circular, quadripartite structure of plastid genomes which includes two inverted repeat regions renders the automatic assembly of plastid genomes challenging. The correct assembly of plastid genomes is a prerequisite for the validity of subsequent analyses on plastid genome structure and evolution. Plastome-based phylogenetic or population genetic investigations, for example, require the precise identification of DNA sequence and length to determine the location of nucleotide polymorphisms. The average coverage depth of a genome assembly is often used as an indicator for assembly quality. Visualizing coverage depth across a draft genome allows users to inspect the quality of the assembly and, where applicable, identify regions of reduced assembly confidence. Based on such visualizations, users can conduct a local re-assembly or other forms of targeted error correction. Few, if any, contemporary software tools can visualize the coverage depth of a plastid genome assembly while taking its quadripartite structure into account, despite the interplay between genome structure and assembly quality. A software tool is needed that visualizes the coverage depth of a plastid genome assembly on a circular, quadripartite map of the plastid genome.

**Results:** We introduce ‘PACVr’, an R package that visualizes the coverage depth of a plastid genome assembly in relation to the circular, quadripartite structure of the genome as well as to the individual plastome genes. The tool allows visualizations on different scales using a variable window approach and also visualizes the equality of gene synteny in the inverted repeat regions of the plastid genome, thus providing an additional measure of assembly quality. As a tool for plastid genomics, PACVr provides the functionality to identify regions of coverage depth above or below user-defined threshold values and helps to identify non-identical IR regions. To allow easy integration into bioinformatic workflows, PACVr can be directly invoked from a Unix shell, thus facilitating its use in automated quality control. We illustrate the application of PACVr on two empirical datasets and compare the resulting visualizations with alternative software tools for displaying plastome sequencing coverage.

**Conclusions:** PACVr provides a user-friendly tool to visualize (a) the coverage depth of a plastid genome assembly on a circular, quadripartite plastome map and in relation to individual plastome genes, and (b) the equality of gene synteny in the inverted repeat regions. It, thus, contributes to optimizing plastid genome assemblies and increasing the reliability of publicly available plastome sequences, especially in light of incongruence among the visualization results of alternative software tools. The software, example datasets, technical documentation, and a tutorial are available with the package at https://github.com/michaelgruenstaeudl/PACVr.

## BACKGROUND

The sequencing and comparison of complete plastid genomes have become a popular method in plant evolutionary research, rendering the precise genome assembly and its quality assessment of high importance. The plastid genomes of most photosynthetically active land plants display a circular, quadripartite structure and comprise two single copy (SC) regions separated by two identical inverted repeats (IR) (Mower and Vickrey, 2018). A total of four partitions with markedly different lengths can, thus, be defined in typical land plant plastomes: the large single copy (LSC) region of ca. 70-90 kilobases (kb), the small single copy (SSC) region of ca. 15-25 kb, and the two IR regions (IRa and IRb) of ca. 20-25 kb each (Ruhlman and Jansen, 2014). The IR regions represent reverse complements of each other and are primarily homogenized through a recombination-mediated replication process (Blazier et al., 2016; Ruhlman et al., 2017). The plastid genomes of most photosynthetically active land plants encode a total of ca. 100-120 proteins, which play a central role in organelle metabolism and photosynthesis (Wicke et al., 2011). Due to their strong structural conservation, uniparental inheritance, a near absence of recombination and a high copy number per plant cell, plastid genomes are highly suitable for comparative genomic studies (Twyford and Ness, 2017). Numerous investigations have sequenced and compared complete plastid genome sequences over the past decade (Gao et al., 2010; Gitzendanner et al., 2018b), and the number of publicly available plastid genomes continues to increase dramatically (Tonti-Filippini et al., 2017). Recent studies on plastid genome structure and evolution now evaluate polymorphisms across hundreds of genome sequences (Bernhardt et al., 2017; Teisher et al., 2017; Saarela et al., 2018; Gitzendanner et al., 2018a), rendering the precise assembly process of plastid genomes and its quality assessment ever more important.

Despite the development of assembly algorithms customized for plastid genomes, the plastome assembly process remains imperfect and often requires the verification, if not manual correction, of the assembly product. Concomitant with the surge in plastid genome sequencing, many new algorithms and pipelines specifically designed for the assembly of plastid genomes have been developed (Ankenbrand et al., 2018; Dierckxsens et al., 2017; Izan et al., 2017; McKain and Wilson, 2017; Coissac, 2017; Gruenstaeudl et al., 2018; Jian et al., 2018; Wang et al., 2018). Most of these tools allow a more accurate and targeted assembly of the plastid genome than generic assembly software, but in many cases some form of manual intervention or post-processing of the assembly results remains necessary (Izan et al., 2017; Gruenstaeudl et al., 2018). The post-processing of automated assembly results often pertains to the correction of the IR length (Tonti-Filippini et al., 2017; Twyford and Ness, 2017), differences in junction boundaries (Wu et al., 2015), and genome circularization (Hunt et al., 2015). Common uncertainties and outright errors in plastid genome assemblies include the inequality of the IR regions in length or sequence (Williams et al., 2015; Gruenstaeudl et al., 2017; Amiryousefi et al., 2018), long homopolymer runs (Kim et al., 2015; Tonti-Filippini et al., 2017), and the imperfect duplication of repeats at the junction of the SC regions (Wu et al., 2015). The differential orientation of the SSC, by contrast, does not constitute an assembly error but reflects the natural presence of heteroplasmy in organellar genomes (Walker et al., 2015). To ensure correctness and reproducibility in plastid genome sequencing and analysis, it is paramount to confirm the validity of plastid genome assemblies (Wu et al., 2015; Tonti-Filippini et al., 2017; Gruenstaeudl et al., 2018). Many of the ambiguities and putative errors recognized in published plastid genome sequences, including those of genome annotations (Tonti-Filippini et al., 2017; Gruenstaeudl et al., 2017; Amiryousefi et al., 2018), could potentially be averted by the application of simple quality assessment strategies (Kim et al., 2015; Gruenstaeudl et al., 2018).

Several measures have been used to indicate the quality of plastid genome assemblies, including contiguity metrics and the length and sequence equality of the IR regions, but sequencing coverage remains one of the most popular proxies for assembly quality. In genome research, the length of the shortest among all those contigs that cover at least 50% of a reference genome is often used as an indicator for the quality of a draft genome (Earl et al., 2011). The closer this length is to the complete length of the reference genome, the more confidence is placed in the completeness and, by extension, the quality of the assembly (Alhakami et al., 2017). This concept is one of several contiguity metrics used to indicate the quality of a genome assembly (e.g., NG50, Earl et al. (2011); NA50 and NGA50, Gurevich et al. (2013)). However, these contiguity metrics are difficult to apply to so far unsequenced organisms due to the requirement of a known reference sequence. Another, more specific measure for validating the quality of genome assemblies constitutes the equality of gene synteny across draft genomes or subsections thereof (Udall and Dawe, 2018). The IRs of a plastid genome, for example, represent recombinogenic isomers and, thus, share the same DNA sequence and gene synteny (Palmer, 1983; Blazier et al., 2016); exceptions to this rule are very rare (Turmel et al., 2017). Equality in length, sequence and gene synteny of the IR regions can, thus, be used as an indicator for the quality of the plastid genome assembly as a whole (Gruenstaeudl et al., 2017). The depth of sequencing coverage (‘coverage depth’ hereafter) represents yet another indicator for the quality of a genome assembly (Pedersen et al., 2017). Average coverage depth is defined as the average number of times each nucleotide of a genome region is represented by aligned reads from a sequence set (Sims et al., 2014); it is a unit-less integer. Coverage depth is an important and highly popular indicator for the quality of a genome assembly in biological research (Nagarajan and Pop, 2013; Ekblom and Wolf, 2014). For plastid genomes, coverage depth is reported almost by default in relation to genome assemblies (Twyford and Ness, 2017; McKain et al., 2018) and has been implemented as a quality metric in several plastome assembly pipelines (Izan et al., 2017; McKain and Wilson, 2017; Coissac, 2017). Information on coverage depth is critical for the assessment of large-scale sequence rearrangements or other structural variation of a genome because a greater coverage depth increases the chance that rearrangement endpoints are captured and confirmed by multiple independent reads (Chen et al., 2009; Sims et al., 2014). In the present investigation, coverage depth, as well as gene synteny of the IR regions, are used as specific indicators for plastid genome assembly quality.

Currently available software tools can generate either unpartitioned plots of plastome coverage depth or quadripartite plastome maps, but the simultaneous, user-friendly visualization of both aspects is presently unsupported. When employing currently available software tools, plant biologists must decide if they wish to visualize either plastome sequencing coverage as unpartitioned, often linear plots or - alternatively - the circular, quadripartite structure of a plastid genome. The assembly pipeline FastPlast (McKain and Wilson, 2017), for example, analyzes coverage depth during run-time and, upon genome assembly, generates a linear coverage plot as part of the pipeline execution (Figure 1a). Similarly, the assembly pipeline IOGA (Bakker et al., 2016) generates linear coverage plots during run-time, allowing users to evaluate the progress of the assembly process during different pipeline iterations (Figure 1b). The assembly pipeline ORG.asm (Coissac, 2017) also estimates average coverage depth during the assembly process but does not visualize this metric. None of these assembly pipelines generates visualizations that account for the circular, quadripartite structure of the plastid genome or for the location of the individual plastome genes. On the other hand, several software tools and web-services exist that visualize complete plastid or bacterial genomes as circular maps. The web-service OrganellarGenomeDraw (OGDRAW; Lohse et al., 2013; Greiner et al., 2019), for example, generates circular maps of plastid and mitochondrial genomes and visualizes gene position and GC content across the genomes. Similarly, the software Circleator (Crabtree et al., 2014) generates circular maps of bacterial genomes and can visualize gene position, GC content, and single nucleotide polymorphism locations in comparison to a reference genome. When co-supplied with text-based configuration instructions and a read mapping file, Circleator can also visualize coverage depth on the circular visualizations (Figure 1c), but the configuration instructions are complex, and unless an intricate, multi-layered visualization procedure is applied, additional genome annotations such as genes are not displayed. The software Circos (Krzywinski et al., 2009) can also be used to generate elaborate visualizations of circular genomes, including plastomes (Shi et al., 2013; Korotkova et al., 2014; Hu et al., 2015), but even more bioinformatic expertise is required to generate the source code underlying these visualizations, which is typically beyond the ability of a normal user in plant biology. Several older software tools and web-services for generating circular genome maps also exist (Sato and Ehira, 2003; Stothard and Wishart, 2005; Conant and Wolfe, 2008; Li et al., 2012; Cheng et al., 2013), but their application in recent research has been minimal, and some of these services have become inaccessible (e.g., Sato and Ehira, 2003; Li et al., 2012; Cheng et al., 2013); inaccessible since at least October 2018). To the best of our knowledge, none of the presently available software tools can visualize the coverage depth of a plastid genome assembly on a circular, quadripartite plastome map while simultaneously displaying the locations of the plastome genes and the location and relative sizes of the single copy and the IR regions as well as the equality of gene synteny in the IR regions, especially in a user-friendly fashion.

**Figure 1.**
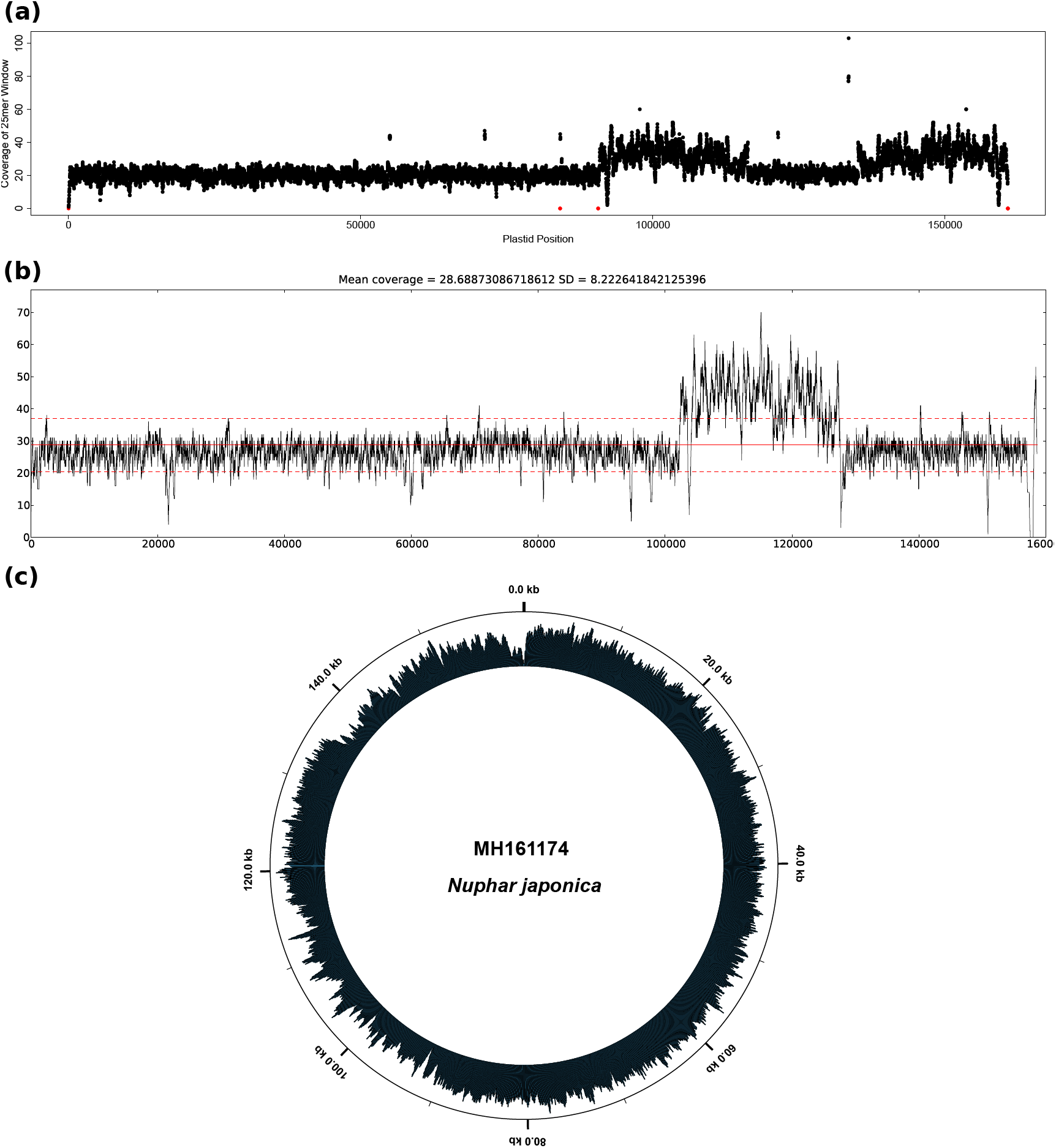
Visualizations of plastome coverage depth by software tools other than PACVr. Displayed is the coverage depth of the plastid genome assembly of *Nuphar japonica* (GenBank accession MH161174) as visualized by the software tools (**a**) FastPlast, (**b**) IOGA, and (**c**) Circleator. For easier viewing, the size of the individual data points was reduced in the visualization by FastPlast, and the tick mark at 160 kb was removed in the visualization by Circleator.

Given the plethora of complete plastid genomes generated in biological research each year (Tonti-Filippini et al., 2017), strong demand for a software tool exists that enables the visual quality assessment of plastid genome assemblies. Specifically, it would be desirable to have a tool that allows users to visually explore the coverage depth of a plastid genome assembly as well as the equality of gene synteny in its IR regions, as both aspects are indicative for the quality of the genome assembly. To be useful to a wide audience, such a software tool must fulfill four criteria: it must (a) be user-friendly and applicable to users with minimal bioinformatics knowledge; (b) generate publication-ready visualizations that allow the determination if and where a genome assembly displays insufficient coverage depth; (c) allow an easy integration into automated workflows or analysis pipelines; and (d) allow users to set customized window sizes and thresholds for coverage depth calculation. Here, we present such a tool, entitled ‘PACVr’ for ‘Plastome Assembly Coverage Visualization in R’. PACVr is a package for the common statistical environment R (R Development Core Team, 2013) that visualizes (i) coverage depth of a plastid genome assembly on a circular, quadripartite plastome map, and (ii) gene synteny between the IR regions of the genome assembly. Specifically, PACVr visualizes coverage depth across the entire plastid genome in user-defined window sizes and in relation to the gene annotations, calculates and displays average coverage depth values for each of the four plastome regions, highlights sectors with coverage depth below a user-specified threshold, and visually connects the gene counterparts of the two IR regions to illustrate equality in gene synteny. By applying PACVr upon plastid genome assembly, users can visually inspect the quality of the assembly and, where applicable, identify regions of reduced confidence or possible errors. Specifically, users can identify sectors of a plastid genome with low coverage depth or missing IR equality and then subject these sectors to error correction or post-processing. Upon presenting the details of this software, we illustrate the application of PACVr on two plastid genome assemblies from different plant families and compare the resulting visualizations with of the output of other software tools for visualizing plastome coverage depth.

## IMPLEMENTATION

### Input and output specifications

The input to PACVr consists of two different files of common file format which contain information on genome sequence and structure as well as coverage depth. Information on genome sequence and structure, as well as the genes encoded by the sequence, is supplied in the GenBank flatfile format. GenBank flatfiles represent the default file type for sequence retrievals from NCBI Nucleotide (Benson et al., 2006) and contain one or more sequence records, with each record comprising general metadata, an annotation table with the names and locations of genes and other sequence features, and the nucleotide sequence itself (Lee et al., 2008). In PACVr, GenBank flatfiles are parsed via the R package genbankr (Becker and Lawrence, 2019) and must, thus, contain only a single sequence record per file, with the locus name no longer than ten alphanumeric characters. Moreover, genbankr requires the location of sequence features that span multiple positions or occur on complementary strands to be specified with the use of only a single invocation of the commands ‘join’ and ‘complement’ each, and all sequence features of class ‘exon’ to be removed. To be suitable for PACVr, the sequence record of the GenBank file must represent a complete, quadripartite plastid genome, with a total sequence length between 100 kb and 200 kb and features annotations for each of the two IRs (with note-qualifiers that have the text values ‘IRa’ and ‘IRb’, respectively). Information on coverage depth is supplied as an input file in the binary alignment/map (BAM) format, which stores alignment and mapping information (Li et al., 2009) and is typically generated during the mapping of sequence reads to a reference sequence by automated read-mappers (Schbath et al., 2012) such as BWA (Li and Durbin, 2009) or Bowtie (Langmead et al., 2009). The BAM file must hereby be indexed and is, thus, accompanied by an ancillary index file. Additional types of information can also be supplied to PACVr upon software initiation, including the window size used for calculating coverage depth, the threshold below which coverage depth is highlighted, and the name of the output file, but they are optional and have well-tested default values set for them.

The output of PACVr is an annotated plastome map in PDF format. Upon initiation, PACVr generates a circular, quadripartite map of the plastid genome in which coverage depth values are displayed as histogram bars, with bars below a predefined threshold highlighted in red and partition-wide average coverage values superimposed as horizontal, green lines. The map also displays the location of all plastome genes, allowing the user to relate areas of low coverage depth to specific genome regions and genes. The map generated by PACVr is saved in PDF format to a user-defined output file.

### Coverage calculation and display

Coverage depth is calculated by PACVr via the application of user-defined window sizes with the software mosdepth (Pedersen and Quinlan, 2017). Window-based coverage calculations have the flexibility of measuring coverage on customized scales, which can be necessary to account for the variability in read length across different Illumina reagent types or sequencing cycle numbers. Using a sorted BAM file plus its ancillary index file as input, mosdepth rapidly infers the coverage of a particular chromosome by tracking all start and end positions of mapped sequence reads and calculating the cumulative sum of their incremented start positions while decrementing the respective end positions (Pedersen and Quinlan, 2017). Based on the results of mosdepth, PACVr infers the average coverage depth for each of the four regions of the plastid genome (i.e., LSC, SSC, IRa and IRb). Both types of coverage depth information are plotted on the plastome map: window-based depth values are displayed in the form of a circular histogram, with the width of each histogram bar equal to the width of the window size, whereas regional coverage averages are displayed as horizontal, green lines superimposed on the histogram bars. PACVr is, thus, different from most other software tools for visualizing coverage depth, which typically display coverage depth as stacked sequence reads (Kearse et al., 2012; Karolchik et al., 2012), line graphs (Bakker et al., 2016), dot graphs (McKain and Wilson, 2017) or bedGraphs plots (Phanstiel et al., 2014), and primarily on linear representations of the input genome.

### IR equality assessment and display

Equality among the IR regions is evaluated by PACVr via the comparison of number, length, and location of all IR genes. Specifically, the software conducts a two-step procedure in which equality in sequence length and number of genes is confirmed across the two IR regions and the equality then visualized by connecting matching genes via blue connector lines. First, PACVr extracts the two IR regions from the input genome, stores the names, the start and the end positions of all IR genes in separate data frames and compares the exact number, length and location of the genes in both regions. Differences in sequence length or gene complement between the two IRs result in warning messages to the user. PACVr then visualizes the equality between the IRs by connecting genes with identical names across the regions using blue lines. The lines hereby originate and end at the central nucleotide of each gene shared between the two IRs. Any difference in name or location of the IR genes becomes visible through unequal or missing connector lines, thus enabling the visual assessment of equality among the IR regions regarding gene presence and synteny. This visualization of gene location and synteny contributes to the discovery of rules and patterns in genome orientation and rearrangements (Wang and Yu, 2014).

### Visualization

PACVr employs RCircos (Zhang et al., 2013) as the visualization engine. RCircos is an R implementation of the Circos environment Krzywinski et al. (2009) and is employed by PACVr to visualize the various aspects of plastome structure and coverage depth in four separate layers. In the first, outermost layer, PACVr plots the names and relative positions of the individual regions of the quadripartite genome structure (i.e., LSC, SSC, IRa and IRb), with each region marked in a different color for easier delineation. In the second layer, PACVr plots the names and positions of all genes of the plastid genome, with gene positions indicated by their central nucleotide. In the third layer, PACVr plots the coverage depth of the plastid genome in the form of a circular histogram, with bars displaying one of two possible colors depending on their depth value relative to a user-defined threshold: bars with a coverage depth above the threshold are displayed in black, bars below that threshold in red. The third layer also displays the average coverage depth of each region via a horizontal, green line, which is missing in areas without coverage. In the fourth, innermost layer, PACVr plots blue lines that connect genes with identical names across the two IR regions, with lines originating and ending at the central nucleotide of each gene. At the top right of the circular graph, PACVr prints a legend that displays the numeric values of average coverage depth of different regions. The name of the organism under study, which is as parsed from the GenBank input file, is displayed as figure title.

### Accounting for quadripartite structure

The quadripartite structure of plastid genomes requires adjustments in the calculation of coverage depth and the visualization of IR equality compared to unpartitioned chromosomes. By default, PACVr calculates window-sized coverage depth values and, based on these, the region-wide average for each of the four plastome regions. However, PACVr would double-count the coverage of those windows that span across a region boundary, unless the coverage calculation included a special adjustment. Similarly, PACVr requires customization when visualizing the equality of gene position and synteny between the two IR regions. Natural expansions in IR size can cause genes located near the border of a single copy and an IR region to be displaced from one region into the other over time (Dugas et al., 2015; Zhu et al., 2016). Without a customized visualization, genes that are located primarily in the single copy region but span into the IR (or vice versa) would not be included in the equality visualization if the central nucleotide of genes used to connect the IR counterparts was located outside the IR. This can be particularly problematic with large genes such as *ycf1* and *ycf2*, which are located near the 5’ end of the SSC and the IRa, respectively, in most angiosperms and represent nearly 10% of the unit-genome length (Ruhlman and Jansen, 2018). A similar issue would arise with trans-spliced genes such as *rps12*, whose exons are located in both the LSC and the IR (Hildebrand et al., 1988). Thus, the code of PACVr was customized to split genes that span more than one genome region into two separate parts along the region boundary, and to treat both parts as separate, gene-like entities. Details on the implementation of these adjustments and their advantages in both coverage calculation and IR equality visualization are described in (Jenke, 2018).

### Installation, dependencies and usage

PACVr was written in R and can be installed via `install github(‘michaelgruenstaeudl/PACVr’`). It requires the presence of the R packages optparse (Davis, 2019), genbankr (Becker and Lawrence, 2019), and RCircos (Zhang et al., 2013) as dependencies; each of these packages is automatically added to the system upon the installation of PACVr. Additionally, PACVr requires the software mos-depth (Pedersen and Quinlan, 2017) to be present on the system, which can be installed via the Unix shell command `conda install mosdepth`. The source code of PACVr is available via Github at https://github.com/michaelgruenstaeudl/PACVr. The technical documentation and a user tutorial (vignette) is distributed as part of the R package.

Two mandatory and five optional input parameters can be specified when invoking PACVr. The mandatory input parameters are: the name of, and file path to, the input GenBank file, and the name of, and file path to, a sorted and indexed BAM file. The optional input parameters are: the window size for calculating coverage depth, with a default value of 250; the coverage depth threshold for the switch in histogram color, with a default value of 25; the command to execute mosdepth, with a default command of `mosdepth;` the decision to store temporary files generated during the coverage depth calculation, with the default set to false; and the name of, and file path to, the output file, with the output saved as `./PACVr output.pdf` by default. The software can be invoked either from within the R environment or directly from a Unix shell. A complete list of the short- and long-flag command-line (CLI) arguments available when invoking PACVr from the Unix shell is displayed via the shell command `Rscript PACVr.R -h / --help`. In the framework of an automated workflow (and upon setting the location of PACVr to a shell variable with the same name), the following shell command can, for example, be used to execute PACVr on one of the empirical datasets co-supplied with the R package:

~~~
Rscript $PACVr/inst/extdata/PACVr_Rscript.R \
    -k $PACVr/inst/extdata/MH161174/MH161174.gb \
    -b $PACVr/inst/extdata/MH161174/MH161174_PlastomeReadsOnly.sorted.bam \
    -r 300 \
    -o MH161174_PlastomeVisualization.pdf
~~~

### Testing of software

To evaluate and demonstrate the functionality of PACVr, the software was tested under a variety of different settings. First, PACVr was tested on empirical data from two different plant families. Specifically, the software was employed for visualizing coverage depth and IR equality of the assemblies of two novel, complete plastid genomes. The plastid genomes represent the species *Dasyphyllum excelsum* (Asteraceae; GenBank accession MH899017) and *Nuphar japonica* (Nymphaeaceae; MH161174). The assemblies of both genomes were generated via Illumina MiSeq sequencing following the sequencing protocol of Gruenstaeudl et al. (2017) and the assembly protocol of Gruenstaeudl et al. (2018). To keep the size of the BAM input files at a maximum of 2.5 megabytes for each test genome and, thus, ensure a lightweight distribution of the R package, the coverage depth of both assemblies was capped at 20 by applying script ‘bbnorm.sh’ of the software BBTools v.33.89 (Bushnell, 2015) on the original sequence reads. This script caps sequence coverage via a stochastic normalization procedure and, thus, results in maximum coverage depths that are approximate. PACVr was employed on both genome assemblies using default parameter values, except for the threshold of coverage depth below which histogram bars are displayed in red, which was set to 15. Second, PACVr was tested on four different operating systems. The operating systems used for testing were: macOS 10.13, Arch Linux 4.18, Debian 9.9 and Ubuntu 18.10. Under each system, PACVr was invoked both from within the R environment as well as directly from a Unix shell. Third, PACVr was compared to alternative software tools for visualizing plastome sequencing coverage. Specifically, we applied three different software tools on the same empirical datasets as employed for testing PACVr itself and then compared their output to the visualization generated by PACVr. The alternative software tools evaluated were: Circleator v.1.0.2, FastPlast v.1.2.8, and IOGA v.20160910.

## RESULTS

### Visualizations by PACVr

PACVr was successfully applied in the visualization of coverage depth and IR equality of two complete plastid genomes used for evaluating the functionality of the software. Specifically, PACVr was successfully employed to visualize the coverage depth of each plastid genome in relation to its circular, quadripartite genome structure and the equality of its IR regions regarding gene position and synteny. For both assemblies, the visualizations indicated genomic areas with markedly lower coverage depth compared to other areas of the same genome, particularly in the IR regions. In the plastid genome of *Nuphar japonica* (Figure 2), the average coverage depth of both IRs was detected to be 16, compared to 19 and 20 for the large and the small SC region, respectively. Moreover, a window-sized coverage depth below the user-selected threshold of 15 was recovered in several locations of the IR, particularly at the 5’ end of the IRa, which corresponds to the location of the ribosomal RNA genes *rrn5* and *rrn4.5*. A suboptimal coverage depth was also detected in one calculation window of the LSC (near *trnV*-TAC, a gene encoding one of the transfer RNAs for valine). In the plastid genome of *Dasyphyllum excelsum* (Additional file 1: Figure S1), the average coverage depth of the IRs was detected to be 17 and 16, respectively, compared to 22 and 21 for the large and the small SC region, respectively. A window-sized coverage depth below the threshold of 15 was found in several locations of the IRs, particularly at the 5’ end of the IRb, which corresponds to the location of gene *ycf2*. A suboptimal coverage depth was also detected in three calculation windows of the LSC and the SSC of the plastid genome of *Dasyphyllum excelsum*. At the same time, no region was identified without sequencing coverage in any of the genome assemblies evaluated, as illustrated by the uninterrupted horizontal, green line across both genome maps. The differences in average coverage depth between the SC and the IR regions may be indicative of a differential assembly quality of these genome partitions, with a lower assembly quality likely present in the IRs. The differences in window-sized coverage depth between the two IR regions, by contrast, are indicative of an unequal or randomized read mapping process during the generation of the input BAM files, as the underlying DNA sequences of the two IR regions are identical in both assemblies. The visualization of equality between the IR regions via blue connector lines indicated equal gene positions and equal gene synteny in both plastid genomes under study. In summary, the IR regions of both plastid genomes evaluated were found to display areas of reduced coverage depth, but sequence length, gene position, and gene synteny in the two IRs were confirmed to be identical. Identical visualizations were retrieved when executing PACVr on macOS and three different Linux distributions, confirming the compatibility of PACVr to different operating systems.

**Figure 2.**
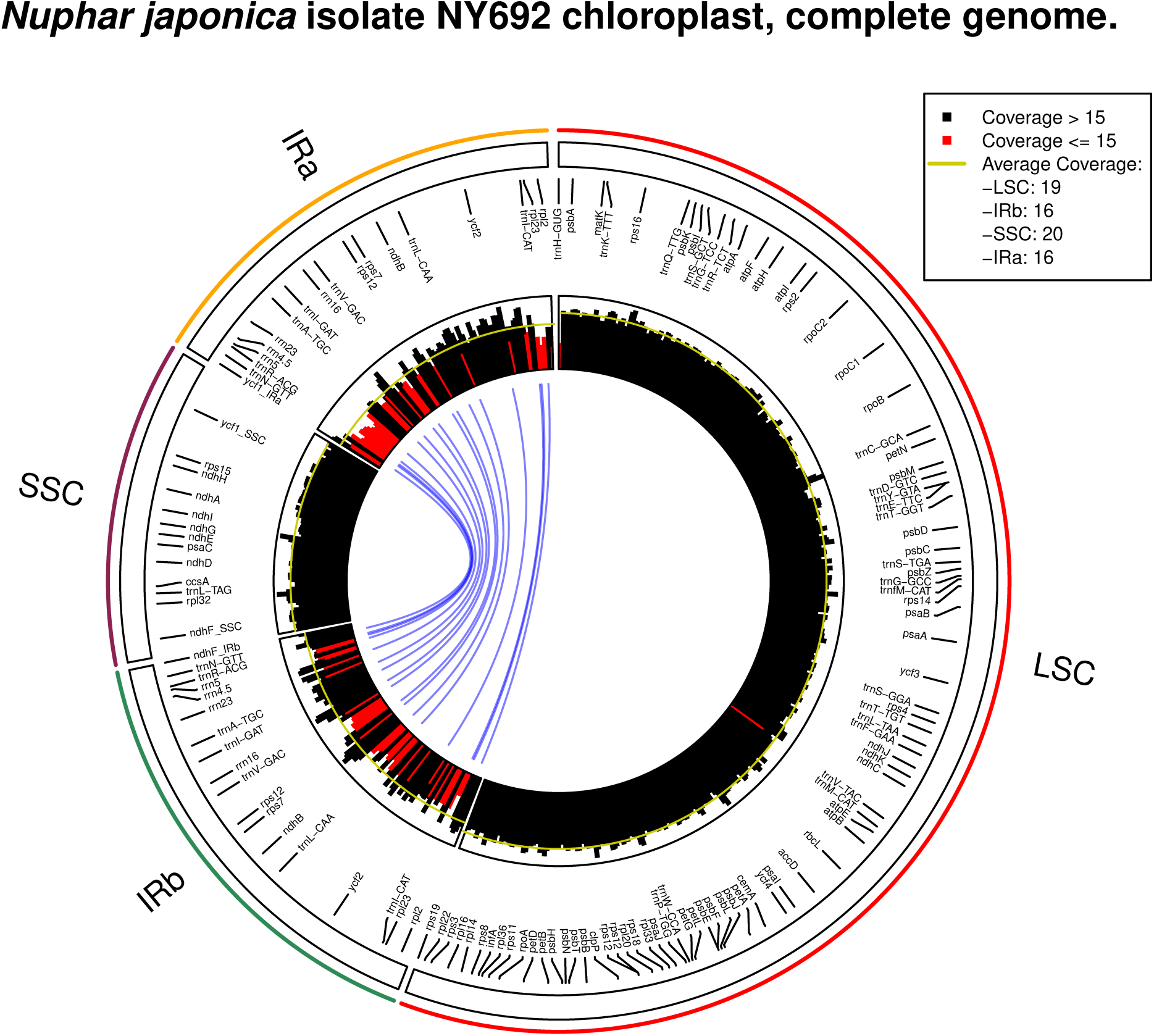
Visualization of coverage depth and IR equality of the plastid genome assembly of *Nuphar japonica* as generated via PACVr.

### Comparison to other software tools

The comparison of visualizations of coverage depth between PACVr and three other software tools recovered incongruent coverage depth distributions, despite applying the same input files. The graphs generated by the tools FastPlast, IOGA and Circleator were incongruent among each other and, in the case of FastPlast and IOGA, also incongruent to PACVr. This incongruence among distributions of coverage depth indicates, among other aspects, that the precise calculation and visualization of coverage depth is insufficiently defined and standardized in the evaluated software tools. For the plastid genome assembly of *Nuphar japonica*, FastPlast generated a linear plot of coverage depth that indicated a higher depth in the IR than in the large and small SC regions (Figure 1a). The coverage depth of the IRs was hereby often larger than 20 and, thus, larger than the manual cap instituted when generating the input files, indicating that the read mapping procedure employed by FastPlast allows the same reads to be counted multiple times across different locations of the input genome. IOGA also generated a linear plot of coverage depth, which indicated a markedly higher coverage depth in an area that approximately corresponds to the SSC compared to other regions of the plastid genome (Figure 1b); the precise location of this area in relation to the overall genome structure is not possible, however, as IOGA generates coverage graphs on the concatenation of individual contigs constructed during the assembly process, and these contigs may not be correctly ordered. Similar to FastPlast, the coverage depth inferred by IOGA surpassed the maximum cap of 20 for certain regions of the genome assembly, indicating a multiple counting of the reads. The graph generated by Circleator was the most similar representation of coverage depth compared to the visualization generated by PACVr. Visualized as a circular plot, it indicated areas of reduced coverage depth in the IRs compared to the SC regions and a largely homogeneous coverage depth across the SC regions (Figure 1c). The precise locations of areas with reduced coverage depth were, however, difficult to determine due to missing references to the quadripartite genome structure and to gene positions. The visualizations of coverage depth for the plastid genome assembly of *Dasyphyllum excelsum* were similarly incongruent among each other as well as to those of PACVr (Additional file 1: Figure S2). In addition, the coverage graph generated by IOGA for this genome displayed coverage for only approximately 130,000 bp of the full genome length due to a missing contig (probably an IR) in the assembly product generated by this software (Additional file 1: Figure S2b). In summary, the visualizations of plastome sequencing coverage with software tools other than PACVr are different in both design and interpretation and can, thus, not be used for comparisons of inferred coverage depth across tools.

## DISCUSSION

### Importance of coverage information in plastid genomics

By visualizing coverage depth in relation to the quadripartite genome structure of plastid genomes and the location of individual genes, PACVr fills the need for a software tool that produces graphically intuitive visualizations for the identification of assembly regions with suboptimal coverage depth. Measuring the coverage depth is critical for the quality assessment of genome assemblies (Pedersen et al., 2017). First, coverage depth is an essential metric for the identification of structural variation, as the depth of sequencing coverage drives the power to detect sequence rearrangements and other structural variants (Sims et al., 2014). Generally, greater coverage depth increases the chance that rearrangement endpoints are captured and confirmed by multiple independent reads (Chen et al., 2009). This can be particularly relevant in the comparison of complete plastid genomes, which often differ structurally (Wang and Yu, 2014), not least in the precise start and end positions of the IR regions (Zhu et al., 2016). Second, coverage depth is an essential metric for the detection of sequence variation, as genomic regions with exceptionally high or low coverage depths become unreliable for variant calling (Li, 2014). In plastid genomics, variant calling can be relevant to identify intra-individual polymorphisms which are typically generated by the effects of heteroplasmy and common to organellar genomes (Scarcelli et al., 2016). Third, de novo assembly algorithms typically operate under the assumption of even coverage depth across the target genome (Peng et al., 2012; Twyford and Ness, 2017), and errors of plastid genome sequences are typically correlated with exceptionally high or low coverage depths (Kim et al., 2015). The visualization of coverage depth of plastome assemblies, thus, represents an important tool in their quality assessment and should be conducted as early in their bioinformatic processing as possible in order to identify problematic assemblies before proceeding with subsequent analyses. Preferentially, such visualizations should be rapid, easily integrable into automated workflows and suitable for the evaluation of a large cohort of genome assemblies (Pedersen et al., 2017).

### Integration into automated pipelines

Given the demand for high throughput in bioinformatic workflows, individual software tools must be easy to integrate into automated analysis pipelines to be of lasting value for the research community. The integration of plastome assembly and annotation into automated or semi-automated workflows has been proposed and conducted by several investigations (McKain et al., 2017; Gruenstaeudl et al., 2018; Jian et al., 2018). Such workflows are designed to deliver more consistent and repeatable results than the manual administration of individual software tools and provide an ideal platform for the integration of assembly quality tests. However, quality management has so far remained unimplemented in most plastid genome analysis pipelines (but see Gruenstaeudl et al. (2018)). In fact, most quality control tools for plastid genome assembly in existence do not provide rapid visualizations of coverage depth. As a result, inaccurate or unsupported plastid genome assemblies may remain undetected and confound subsequent analyses, especially in large, composite investigations that compare hundreds if not thousands of plastid genomes (e.g., Saarela et al., 2018; Gitzendanner et al., 2018a). Hence, it is critical to visualize the coverage profile of a plastid genome through an automated, yet user-friendly process. Strong emphasis was, thus, placed on the ability to easily integrate PACVr into automated bioinformatic pipelines. Given this objective, PACVr was tested on all major operating systems. Similarly, PACVr was designed to enable an operation directly from the Unix shell using CLI arguments, which allows easy integration into automated workflows.

### Importance of open-source software in plastid genomics

Several previously available web-services for visualizing circular plastome maps have become inaccessible over recent years, highlighting the importance of open-source software development in plastid genomics. The development and release of PACVr as an open-source software tool was one of the guiding principles in its development, as this allows other researchers to independently access its source code, customize the software, and extend its functionality. The aim for open-source development is particularly important in the field of plastid genomics, where several previously developed web-services have become inaccessible over recent years. In fact, several interactive web-based tools had been developed to visualize circular chromosomes and their associated metadata, including complete plastid genomes. However, many of these tools are no longer applied because their online interfaces have lost connectivity to the world wide web, and their source code has never been made publicly available. The online platform CARAS (Li et al., 2012), for example, offered functionality to annotate and visualize complete plastid genomes and save the results in different output formats, but its web service has been inaccessible since at least February 2017. Similarly, the web platform CGAP (Cheng et al., 2013) offered functionality to generate circular or linear genome maps, annotate assembled plastid genomes and conduct comparative plastome analysis, but has been inaccessible since at least April 2017. Some of these online services provided installation-free alternatives to the limited number of visualization software tools for plastid genomics (Sablok et al., 2016), and their inaccessibility should be considered a loss for the plant biological research community. Had these services been developed as open-source projects, other researchers would have had the opportunity to continue maintenance and development of these resources (Huang et al., 2017). Open-source development and public accessibility of software tools are, thus, considered critical aspects of bioinformatic software development (Ince et al., 2012; Howison and Bullard, 2016; Darriba et al., 2018). Consequently, PACVr was developed as an open-source R package that is publicly available via GitHub.

## CONCLUSIONS

Coverage depth is often used as an indicator for the quality of a plastid genome assembly. The R package PACVr was designed to visualize coverage depth of plastome assemblies in relation to the circular, quadripartite structure of plastid genomes, the location of individual plastome genes, different window calculation sizes and user-defined threshold values for coverage depth. PACVr also enables the visual assessment of equality among the IR regions regarding gene presence and synteny. In tests on empirical data, the software successfully visualized the coverage depth and IR equality of complete plastid genomes of different plant families. Its evaluation also highlighted that alternative coverage visualization tools for plastid genomes generate incongruent depth visualizations on the same input data. Given its design as an open-source R package with a Unix shell interface, PACVr allows easy integration into bioinformatic pipelines and, thus, provides an important tool for automated quality control in plastid genome sequencing.

## Acknowledgments

The authors thank Yannick Hartmaring of the Freie Universität Berlin for assistance with testing the final software version. The authors acknowledge the high-performance computing service of the ZEDAT of the Freie Universität Berlin for providing allocations of computing time. The development of code for of this R package constitutes part of a thesis by NJ toward a bachelor of science degree.

## Authors’ Contributions

MG - Conception, project design, oversight, implementation, documentation, testing on empirical data, manuscript writing and manuscript review. NJ - implementation, documentation and testing. Both authors have read and approved the final version of the manuscript.

